# Transplantation of Muscle Stem Cell Mitochondria Rejuvenates the Bioenergetic Function of Dystrophic Muscle

**DOI:** 10.1101/2020.04.17.017822

**Authors:** Mahir Mohiuddin, Jeongmoon J. Choi, Nan Hee Lee, Hyeonsoo Jeong, Shannon E. Anderson, Woojin M. Han, Berna Aliya, Tsvetomira Z. Peykova, Sumit Verma, Andrés J. García, Carlos A. Aguilar, Young C. Jang

## Abstract

Mitochondrial dysfunction has been implicated in various pathologies, including muscular dystrophies. During muscle regeneration, resident stem cells, also known as muscle satellite cells (MuSCs), undergo myogenic differentiation to form *de novo* myofibers or fuse to existing syncytia. Leveraging this cell-cell fusion process, we postulated that mitochondria stemming from MuSCs could be transferred to myofibers during muscle regeneration to remodel the mitochondrial network and restore bioenergetic function. Here, we report that dystrophic MuSCs manifest significant mitochondrial dysfunction and fuse with existing dystrophic myofibers to propagate mitochondrial dysfunction during muscle repair. We demonstrate that by transplanting healthy donor MuSCs into dystrophic host muscle, the mitochondrial network (reticulum) and bioenergetic function can be rejuvenated. Conversely, when bioenergetically-compromised donor MuSCs are transplanted, improvements in mitochondrial organization and bioenergetic function were ablated in the dystrophic recipient. Overall, these data reveal a unique role of muscle stem cells as an essential regulator of myofiber mitochondrial homeostasis and a potential therapeutic target against mitochondrial myopathies.

## Introduction

Duchenne muscular dystrophy (DMD) is a debilitating and fatal skeletal muscle disorder characterized by a loss-of-function mutation in dystrophin, a structural protein that connects the membrane and cytoskeleton of the myofiber to its surrounding extracellular matrix (Bonilla et al., 1988; Campbell, 1995; Hoffman et al., 1987). As a consequence, unstable sarcolemmal membranes (Hamer et al., 2002) cause dystrophic myofibers to become susceptible to contraction-induced injuries, invoking progressive muscle wasting (Schmalbruch, 1984). To repair damaged muscle, quiescent muscle stem cells (MuSCs), also referred to as satellite cells, commit to the myogenic lineage and initiate adult myogenesis through activation of distinct transcriptional factors, followed by differentiation and fusion into *de novo* syncytia or existing multinucleated myofibers (Charge and Rudnicki, 2004). Proliferating MuSCs can also self-renew and replenish the quiescent stem cell pool. However, in DMD, repeated rounds of degeneration and regeneration impair the myogenic progression of MuSCs and eventually deplete the muscle-forming stem cell pool due to the aberrant microenvironment (Dumont and Rudnicki, 2016; Tabebordbar et al., 2013). These pathological features exacerbate the regenerative capacity of DMD myofibers and ultimately lead to loss of ambulatory function and early death caused by the failure of vital respiratory muscles (Blau et al., 1983). Hence, considerable research effort has been directed towards developing cell-based therapies to restore dystrophin and regenerative function in pre-clinical and clinical models of DMD (Sun et al., 2020).

Growing evidence suggests that the loss of dystrophin is accompanied by myofiber mitochondrial dysfunction and a concomitant increase in oxidative stress (Hughes et al., 2019; Kuznetsov et al., 1998; Pant et al., 2015; Schuh et al., 2012; Timpani et al., 2015b). Moreover, recent studies reported that restoration of mitochondrial bioenergetic function ameliorates pathological progression in the mouse model of DMD (Ryu et al., 2016; Zhang et al., 2016). Several investigators have reported correlative evidence that the unstable sarcolemmal membrane and contraction-induced injuries cause an overwhelming influx of extracellular Ca^2+^ that alters buffering capacity and this, in turn, leads to a loss of mitochondrial membrane potential, a decline in oxidative phosphorylation, an increase in reactive oxygen species (ROS) production, and apoptotic or necrotic death of myofibers (Franco and Lansman, 1990; Ryu et al., 2016; Timpani et al., 2015b; Vila et al., 2017). Recently, we demonstrated that during muscle regeneration, damaged mitochondria are removed by macro-autophagy (mitophagy), and newly synthesized mitochondria stemming from satellite cells reconstitute mitochondrial networks to reestablish mitochondrial-nuclear genome communication in regenerating myofibers (Mohiuddin et al., 2019). As myofibers are both multinucleated and post-mitotic, the relationship between myonuclei and the mitochondrial network is particularly important in mature skeletal muscle. Approximately 99% of mitochondrial proteins are nuclear-encoded and must be transported from myonuclei into mitochondria (Ryan and Hoogenraad, 2007). Thus, without competent accretion of new myonuclei from MuSCs and a proper formation of the myonuclear domains, the cytoplasmic volumes in which each nucleus governs, mitochondrial homeostasis and bioenergetic function are severely compromised (Jang et al., 2010; Murach et al., 2018; Qaisar and Larsson, 2014; Rosser et al., 2002). In addition, muscle mitochondria form myofibrillar and subsarcolemmal networks, allowing an efficient exchange of nascent proteins encoded by different myonuclei (Glancy et al., 2015). Furthermore, despite their confined locations within myofibers, muscle mitochondria readily regulate their size, shape, and volume through mitochondrial dynamics (fission, fusion, and mitophagy) to maintain plasticity (Casuso and Huertas, 2020; Glancy et al., 2018; Mishra et al., 2015). Yet, very little is known on whether recurring degeneration and impaired asymmetric division of MuSCs impact the formation of the mitochondrial network and whether putative mitochondrial dysfunction found in DMD myofibers arise from MuSCs as a consequence of repetitive rounds of adult myogenesis.

Herein, we demonstrate that MuSC mitochondria play a causal role in the bioenergetic function of mature myofibers. In dystrophin-deficient MuSCs, which embody defective mitochondria, bioenergetically-compromised mitochondria are transferred to myofibers during muscle regeneration. By leveraging ectopic MuSC transplantation, we show that healthy donor MuSCs can reconstitute the mitochondrial network of dystrophic host muscle and ameliorate mitochondrial function. Conversely, the transplantation of MuSCs with damaged mitochondria abolishes metabolic enrichment, verifying that the quality of donor MuSC mitochondria is critical in boosting the oxidative capacity of the diseased muscle during muscle regeneration. These findings provide a conceptual foundation for using MuSC-derived mitochondrial transplantation as a therapeutic tool that can be applied to numerous skeletal muscle disorders, including mitochondrial myopathies.

## Results

### MuSCs from dystrophin-deficient mdx mice exhibit mitochondrial dysfunction

Studies have reported that muscles from *mdx* mice, an animal model of Duchenne muscular dystrophy (DMD), show significant deficits in mitochondrial function (Bell et al., 2019; Kuznetsov et al., 1998; Percival et al., 2013; Ryu et al., 2016; Schuh et al., 2012). To determine whether mitochondrial dysfunction in *mdx* myofibers originate from MuSCs, FACS-purified quiescent MuSCs from young wildtype and *mdx* mice were compared. First, to examine morphological differences, mitochondrial reporter (*mitoDendra2*) mice, in which the fluorescent protein *Dendra2* is targeted to the mitochondrial matrix (Pham et al., 2012), were crossed with *mdx* mice and their MuSCs were compared to MuSCs isolated from *mitoDendra2* (referred to as WT). Whereas overall MuSC frequency was not different, the distribution of mitochondrial volume per MuSC was towards greater volume in quiescent muscle stem cells from *mdx* mice compared to WT controls (Figure 1A, 1B, S1A, and S1B). Next, to quantify differences in mitochondrial molecular signature and metabolism, we assessed bulk RNA sequencing on freshly isolated MuSCs from WT and *mdx* mice. The transcriptome profile revealed that gene sets that regulate mitochondrial protein complex and mitochondrial inner membrane were among the most differentially enriched pathways in quiescent *mdx* MuSCs compared to WT control cells (Figures 1C, S2A, S2B). Specifically, in *mdx* MuSCs, the majority of the nuclear-encoded mitochondrial electron transport chain (ETC) genes were upregulated, supporting the notion that dystrophic MuSCs comprise of higher mitochondrial content than WT MuSCs. In stark contrast, all 13 ETC genes encoded by mitochondrial DNA were dramatically downregulated in *mdx* MuSCs compared to WT MuSCs, suggesting a potential disruption in nuclear and mitochondrial genome communication.

**Figure 1.**
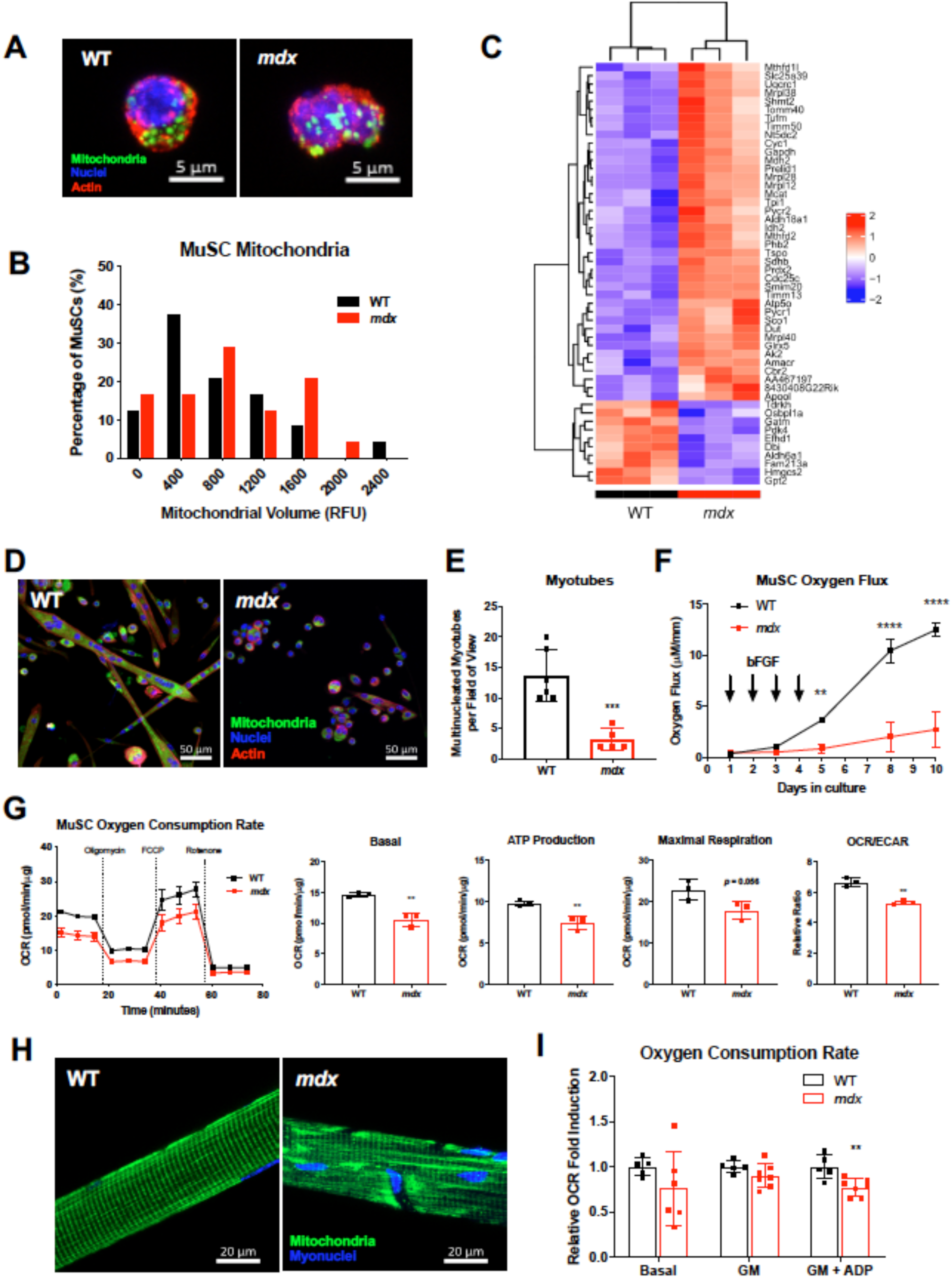
Mitochondrial dysfunction in *mdx* muscle stem cells. (A) Representative confocal images of quiescent MuSCs purified from wild-type (WT)/mitoDendra2 and *mdx*/mitoDendra2. Mitochondria are depicted in green, nuclei in blue, F-actin in red. (B) Histogram of mitochondrial content (relative fluorescent units) of 25 MuSCs. (C) Hierarchical clustering and heatmap of top fifty differentially regulated mitochondrial gene expression (D) Representative confocal images of MuSC-derived myotubes from WT/mitoDendra2 and *mdx*/mitoDendra2 mice. Mitochondria in green, nuclei in blue, actin in red. (E) Quantification of multinucleated myotubes within each field of view. Each sample represents biological replicates (n=6). (F) Comparison of muscle stem cell oxygen flux rate *ex vivo*. FACS purified MuSCs from WT and *mdx* were monitored for 10 days. (G) Oxygen consumption rates (OCR), ATP production, and OCR/extracellular acidification rate (ECAR) of MuSCs cultured for 8 days with daily 25 ng/ml bFGF supplementation using a Seahorse XFp Analyzer (n=3). (H) Representative confocal images of isolated single myofiber from WT/mitoDendra2 and *mdx*/mitoDendra2 tibialis anterior muscle. Mitochondria in green, myonuclei in blue. (I) Bioenergetic functional measurement of permeabilized myofibers from wildtype and *mdx* tibialis anterior muscle. Basal (no substrate), state 2 (Complex I-linked substrate, glutamate and malate (GM), and state 3 (GM+ADP) respiration of (n≥5). Mean ± SD \*\**p*<0.01, \*\*\**p*<0.001, \*\*\*\**p*<0.0001 compared to wildtype for all figures.

It has been shown that MuSCs from *mdx* exhibit significant deficits in myogenesis *ex vivo* (Dumont et al., 2015a; Xu et al., 2013). Moreover, dystrophic muscle *in vivo* partially loses the MuSC subpopulation that co-expresses low levels of β1-integrin and CXCR4, which are required to maintain quiescence (Dumont et al., 2015b; Rozo et al., 2016), suggesting an abnormal balance between self-renewing and committed MuSCs (Figure S1C). To test whether the altered myogenic capability is linked to mitochondrial bioenergetic function, kinetic oxygen flux measurements, a metric for oxidative respiration, were collected from cultured MuSCs through myogenic activation, proliferation, differentiation into myotubes. Although WT MuSCs show a remarkable increase in respiration to cope with the high energy demands of myotube formation (Cerletti et al., 2012; Mohiuddin et al., 2019; Ryall, 2013), the ability of *mdx* MuSCs to fuse into myotubes was significantly compromised (Figure 1D, 1E), and oxidative capacity was attenuated throughout myogenesis (Figure 1F). To corroborate that the reduction in bioenergetics seen in *mdx* MuSCs is independent of myotube content, we assessed the oxygen flux using mitochondrial ETC inhibitors and normalized the data to total protein content. The basal oxygen consumption rate (OCR) of dystrophic MuSCs was reduced compared to WT cells, even when normalized to total protein (Figure 1G). When MuSC mitochondria were stimulated with mitochondrial ETC inhibitors, *mdx* MuSCs also exhibited deficiencies in ATP production and maximal respiration (Figures 1G, S1D), showing impaired mitochondrial efficiency and responsiveness to the energy demands of myogenic progression.

During muscle regeneration, MuSC-derived myotubes form mature myofibers by fusing their intracellular components, including mitochondria. Thus, we analyzed the mitochondrial morphology of mature myofibers isolated from mitoDendra2 control (WT) and *mdx*/mitoDendra2 (*mdx*) mice. Since bioenergetic function relies on the mitochondrial morphology within the cell (Rafelski, 2013), we first assessed the mitochondrial organization of the myofibers. Whereas myofibers from control mice display a healthy mitochondrial network that is segregated into compartmentalized reticula along the contractile apparatus (Mishra et al., 2015), dystrophic myofibers portrayed a highly disordered mitochondrial morphology (Figures 1H), particularly near the subsarcolemmal region of the myofiber which may be more susceptible to contraction-induced membrane instability (Vila et al., 2017). Next, using permeabilized myofiber bundles loaded into a high-resolution respirometer (Pesta and Gnaiger, 2012), we measured the oxygen consumption rate of myofibers normalized to wet weight at basal respiration without exogenous substrates (state 1), mitochondrial ETC complex I-linked respiration with the addition of glutamate and malate (state 2) (Gnaiger, 2009), and ATP-converting respiration by addition of ADP (state 3). Although there was no statistical difference in OCR between WT and *mdx* myofibers at state 1 or state 2 respiration, state 3 respiration of *mdx* myofibers was significantly reduced (Figure 1I), indicating disrupted bioenergetic function of dystrophic muscle, consistent with previous studies (Kuznetsov et al., 1998). Taken together, these data indicate that dystrophic MuSCs contain defective mitochondria that limit their myogenic capacity and contribute to the mitochondrial dysfunction of dystrophic muscle following fusion into the myofiber, highlighting the MuSC as a potential source of mitochondrial dysfunction in pathological DMD muscle.

### MuSC mitochondrial transplantation

We recently demonstrated that myofiber mitochondrial and myonuclear domains are remodeled by MuSCs during muscle regeneration (Mohiuddin et al., 2019). Since dystrophin-deficient MuSCs embody defective mitochondria that convey to the mature myofiber through myogenic fusion, we interrogated the ability of healthy, transplanted MuSCs to remodel the mitochondria of dystrophic myofibers, generate myonuclei that effectively regulate mitochondrial biogenesis, and improve the bioenergetic function of the myofibers. Because mitochondria are located in the cytoplasm of the cell (Shaw et al., 2008), and the vast majority of mitochondrial proteins (∼99%) are nuclear-encoded (Boengler et al., 2011), we first transplanted purified quiescent MuSCs from transgenic reporter mice with fluorescent proteins expressed in either cytosolic or nuclear compartments into *mdx* muscle to delineate successful transfer of MuSC-derived cytosol and myonuclei to dystrophic myofibers (Figure S3A, S3B). Interestingly, centrally located myonuclei, a spatial characteristic of myonuclei within regenerating myofibers, expressed the nuclear fluorescence marker of transplanted MuSCs, suggesting that these myonuclei were derived from the transplanted MuSC. Next, to visualize MuSC-driven transfer of mitochondria into dystrophic myofibers, we transplanted MuSCs isolated from mitochondrial reporter mice, *mitoDendra2*, into *mdx* recipients (Figure 2A) and indeed found that transplanted MuSC mitochondria are engrafted in recipient myofibers (Figure 2B) and whole muscle homogenate (Figure 2C). Whereas most of the transplanted mitochondria formed an organized interfibrillar mitochondrial network at 28 days following transplantation, we also observed high density of mitochondria in between newly regenerating fibers demarked by centrally located myonuclei (Figure 2B, *bottom*). This implies that MuSC-derived myonuclei and newly synthesized mitochondria form a nuclear-mitochondrial genome communication and integrate into the existing mitochondrial reticula of the recipient muscle. To further validate efficient mitochondrial transfer, we then measured relative mtDNA copy number of muscle homogenates from *mdx* control and transplanted *mdx* muscle. As expected, we observed an elevated mtDNA copy number in transplanted muscle, corroborating the MuSC-driven mitochondrial biogenesis in the recipient muscle (Figure 2D). Furthermore, following mitochondrial transplantation, the overall mitochondrial gene expression drastically upregulated in the recipient muscle (Figure 2E). Among differentially regulated mitochondrial genes in the transplanted muscle, the Slc25 family of proteins showed the most striking changes (Figure 2E). Slc25 is a family of nuclear-encoded mitochondrial proteins that are responsible for transporting substrates and metabolites involved in the TCA cycle into the mitochondria (Palmieri, 2013). We verified this finding of increased mitochondrial content by measuring the protein content of a key mitochondrial electron transport chain component, complex I (NDUFB8 subunit) in MuSC transplanted muscles (Figure 2F). Altogether, these data indicate that donor MuSCs transfer functional mitochondria to the host myofibers, and an increase in healthy mitochondria could lead to bioenergetic enhancements following transplantation.

**Figure 2.**
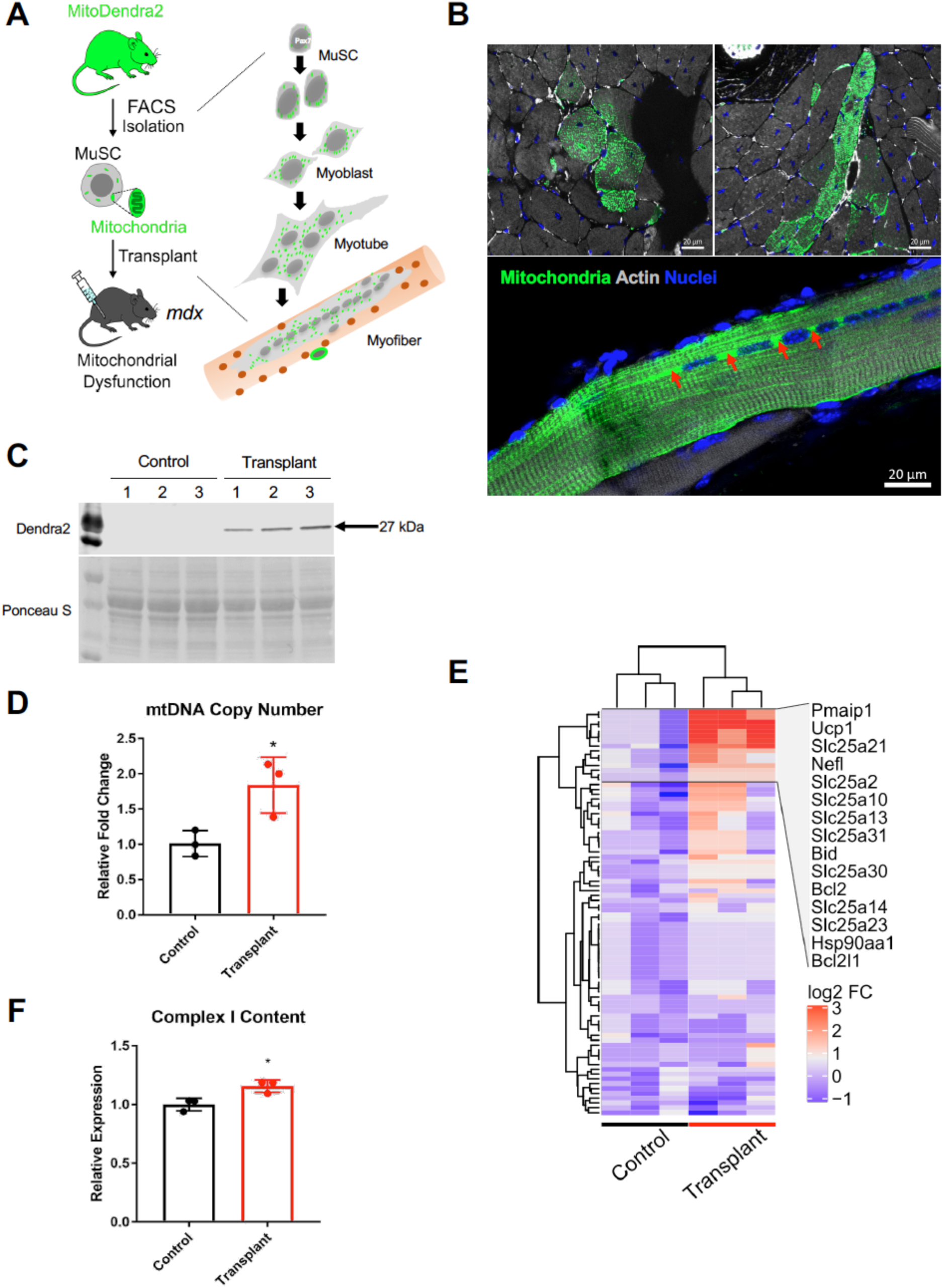
Reconstitution of functional mitochondria via muscle stem cell transplantation. (A) Schematic diagram of mitoDendra2 MuSC transplantation into *mdx* tibialis anterior (TA) muscle. (B) Representative images of mitochondrial transfer. Cross-section of TA and isolated myofiber from TA 28 days following transplantation. Mitochondria in green, actin in gray, myonuclei in blue. Red arrows depict muscle stem cell-derived mitochondria (C) Dendra2 Western blot of muscle homogenate from *mdx* control and MuSC transplanted *mdx* muscle (n=3). (D) mtDNA copy number of *mdx* control and MuSC transplanted TA of *mdx* mice (n=3). (E) Hierarchical clustering of mitochondrial gene expression from *mdx* control and MuSC transplanted *mdx* muscle using PCR array (n=3). (F) Complex I (NDUFB8-subunit) Western blot analysis of muscle homogenate from *mdx* control and MuSC transplanted *mdx* muscle (n=3). Mean ± SD, \**p*<0.05 compared to contralateral control for all figures.

### MuSC-derived mitochondrial transplantation improves bioenergetic function

In the next set of experiments, we examined whether enhanced mitochondrial content via healthy MuSC mitochondrial transplantation resulted in functional improvements of the dystrophic host muscle. To address this, we assessed the morphology and bioenergetic function of the transplanted mitochondria relative to the endogenous *mdx* mitochondria. In multinucleated skeletal muscle, mitochondrion form reticula in distinct locations (Glancy et al., 2018; Glancy et al., 2015; Mishra et al., 2015). The morphological arrangement of mitochondrial reticula is tightly correlated to the cell’s ability to generate energy (Benard et al., 2007; Glancy et al., 2015), so we first compared the various spatial configurations of myofiber mitochondria. Whereas the segregated architecture of endogenous mitochondrial reticula was lost in *mdx/mitoDendra2* myofibers, transplanted MuSC mitochondria from young *mitoDendra2* mice into standard *mdx* mice restored the structural integrity of the mitochondrial network, similar to that of WT/*mitoDendra2* myofibers, and centrally located myonuclei were surrounded by high densities of exogenous mitochondria (Figures 3A, 3B). In support of the high perinuclear mitochondrial density of transplanted muscles, gene expression of PGC-1α, a master regulator of mitochondrial biogenesis (Jager et al., 2007; Ventura-Clapier et al., 2008), was upregulated (Figure 3C), demonstrating that MuSC transplantation remodels the mitochondrial network within the dystrophic myofiber.

**Figure 3.**
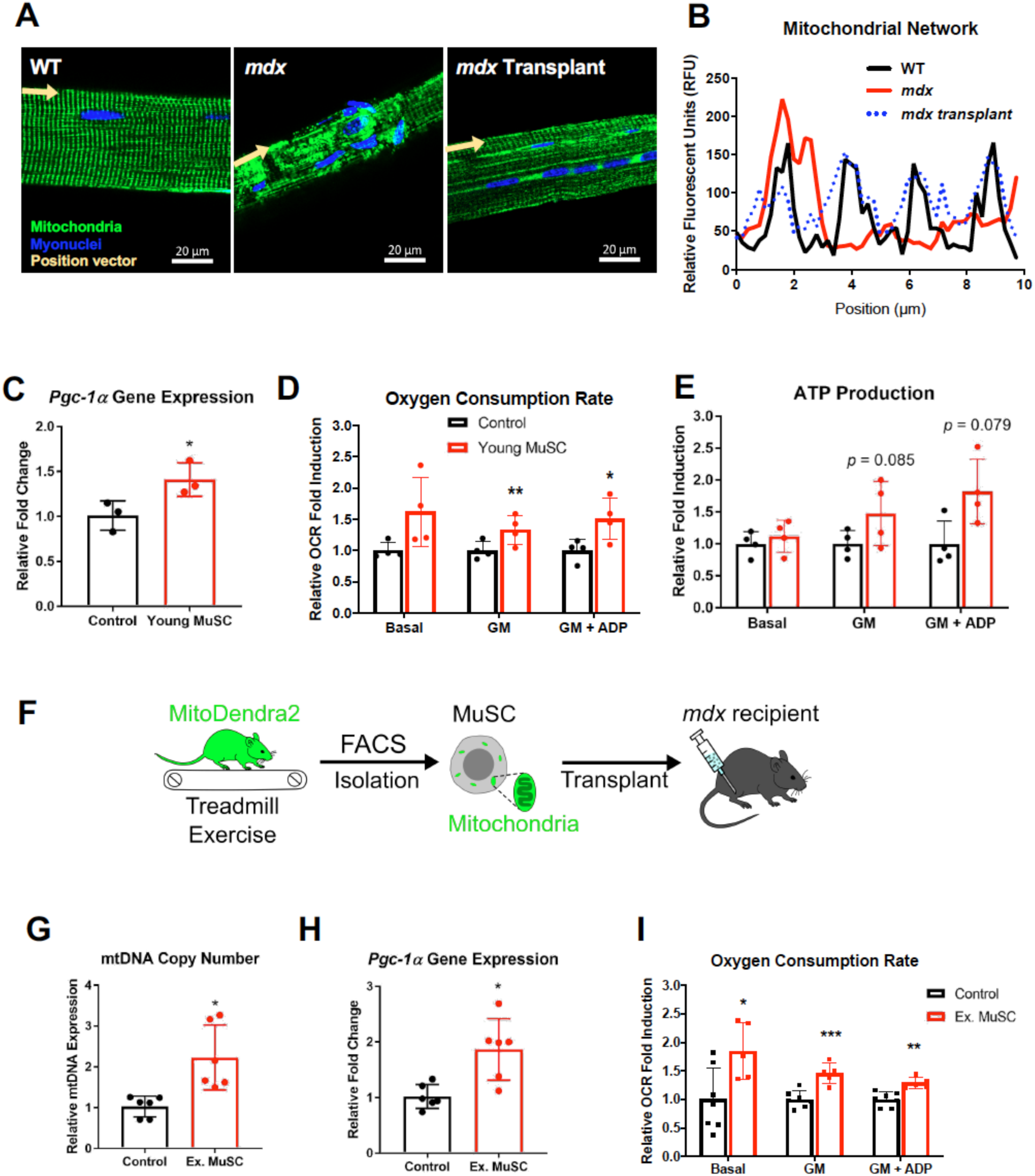
Muscle stem cell mitochondrial transplantation improves bioenergetic function in *mdx* mice. (A) Representative z-stacked images of single fibers from TA of WT/mitoDendra2, *mdx*/mitoDendra2, and *mdx* transplanted with young mitoDendra2 MuSCs. (B) Quantification of mitochondrial network organization (relative fluorescent units) along position vectors (10 µm of fiber length) to portray segregation of reticula. (C) *Pgc-1α* gene expression of contralateral *mdx* control and young MuSC transplanted TA in *mdx* (n=3). (D) States 1, 2, and 3 OCR of permeabilize myofibers from *mdx* control and young MuSC transplanted TA in *mdx* (n=3). (E) ATP production in states 1, 2, and 3 respiration of *mdx* control and young MuSC transplanted TA in *mdx* (n=3). (F) Schematic of exercise-trained donor MuSCs transplanted into *mdx* host muscle. (G) mtDNA copy number of *mdx* control and exercised MuSC transplanted TA in *mdx* (n=6). (H) *Pgc-1α* gene expression of *mdx* control and exercised MuSC transplanted TA in *mdx* (n=6). (I) States 1, 2, and 3 OCR of *mdx* control and exercised MuSC transplanted TA in *mdx* (n=6). Mean ± SD, \**p*<0.05, \*\**p*<0.01, \*\*\**p*<0.001 compared to contralateral control for all figures.

Because mitochondria must form an organized network in order to efficiently distribute ATP to the regenerating myofibers (Wagatsuma et al., 2011), we next performed direct functional analyses of MuSC-derived mitochondria following transplantation to ascertain whether the transferred mitochondria could improve the oxidative phosphorylation activity of dystrophic myofibers. Although there was no statistical difference in oxygen consumption rate between *mdx* controls and transplanted myofibers with their own endogenous substrates at basal respiration, the addition of complex I-linked substrate, glutamate and malate (state 2), as well as supplementation of ADP (state 3) showed significant improvements in oxidative phosphorylation of the engrafted myofibers (Figure 3D). ATP measurements from transplanted myofibers exhibited trends towards increased ATP production at state 2 and state 3 respiration but did not reach statistical significance (Figure 3E). In contrast, when *mdx* donor MuSCs were transplanted into *mdx* recipient muscle, no improvements in mitochondrial biogenesis or oxidative capacity were observed (Figure S4), reaffirming our results of healthy MuSC-derived mitochondrial transplantation driving rejuvenation of the host muscle.

The beneficial effects of endurance exercise training on skeletal muscle mitochondrial biogenesis and mitochondrial function have been well characterized (Baar et al., 2002; Dohm et al., 1973; Hood, 2009; Perry and Hawley, 2018). Likewise, endurance exercise has been shown to promote myogenesis and myonuclear accretion (Hawke, 2005; Shefer et al., 2010; Smith et al., 2001). Although the transplantation efficiency of exercised MuSC mitochondria has not yet been studied, we postulated that transplantation of the bioenergetically-enriched mitochondria from exercise-trained donor MuSCs would propagate to the dystrophic myofibers during myogenesis. Indeed, when we transplanted 8-week endurance-exercised MuSCs as donors (Figure 3F), we observed significantly upregulated mtDNA copy number and PGC-1α expression compared to non-exercised control MuSCs (Figures 3F, 3G, and 3H), implying increased mitochondrial biogenesis in these muscles. To test whether this increased mitochondrial content was associated with improved mitochondrial function, exercised-MuSC-fused *mdx* myofibers were tested in a respirometer. We observed an elevated basal oxygen consumption rate compared to *mdx* control (Fig. 3I), substantiating the increased mitochondrial content demonstrated by mtDNA copy number data. Indicative of improved oxidative function, these transplanted mitochondria also manifested higher state 2 and state 3 oxygen consumption rate than *mdx* control (Fig. 3I), demonstrating that transplanted MuSCs from endurance-exercised MuSCs fuse their mitochondria to dystrophic myofibers to promote mitochondrial biogenesis and restore bioenergetic function. These data reinforce the concept that transplantation of healthy MuSC mitochondria restores not only dystrophin content and myogenic function (Gussoni et al., 1999) but also augments the mitochondrial function of dystrophin-deficient DMD muscle.

### Transplantation of dysfunctional MuSCs does not improve bioenergetic function

To verify that the functional restoration of mitochondria following transplantation depends on the fusion of healthy donor MuSC-derived mitochondria, we performed a complementary analysis by transplanting MuSCs with defective mitochondria and assessing their effect on the bioenergetics of engrafted myofibers. Mitochondrial dysfunction is a hallmark of aging (Figure 4A) that has been thoroughly studied and linked to a number of age-related diseases (Huang and Hood, 2009; Jang et al., 2010; Marzetti et al., 2013; Short et al., 2005). As anticipated, transplanted mitochondria from aged MuSCs (Figure 4B) showed diminished functional improvements following transplantation. Following aged MuSC transplantation, no differences were observed in mtDNA copy number or PGC-1α expression compared to *mdx* control (Figure 4C, 4D), showing that transplantation of aged MuSCs does not increase mitochondrial content. Moreover, the oxygen consumption rate on transplanted myofibers following transplantation of aged MuSCs did not show any signs of improvement (Figure 4E), demonstrating that MuSC-based transplantation of impaired mitochondria does not enhance the bioenergetic function of dystrophic myofibers.

**Figure 4.**
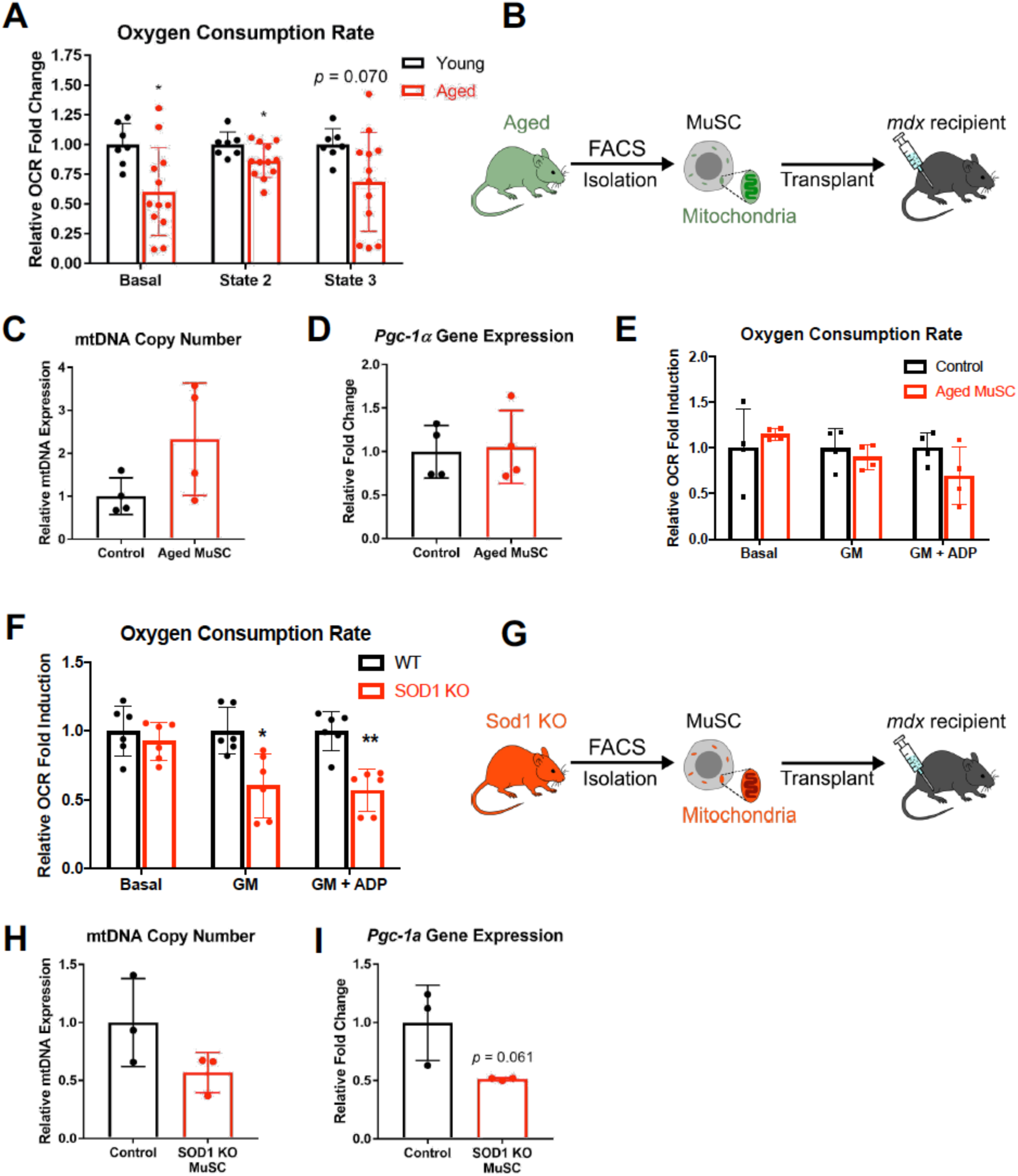
Transplantation of dysfunctional muscle stem cell mitochondria exacerbates mitochondrial dysfunction in *mdx* mice. (A) OCR of permeabilized fibers from the TA of young and aged mice (n≥7). (B) Schematic of aged donor MuSC transplantation into *mdx* host. (C) mtDNA copy number following transplantation of aged MuSCs into *mdx* (n=4). (D) *Pgc-1α* gene expression following transplantation of aged MuSC donors into *mdx* recipients (n=4). (E) States 1, 2, and 3 OCR of permeabilized fibers following transplantation of aged MuSCs into *mdx* host muscle (n=4). (F) OCR of permeabilized fibers from TA of wildtype and *Sod1* knockout mice (n=6). (G) Schematic diagram of *Sod1* KO MuSC donor transplantation into *mdx* recipients. (H) mtDNA copy number following transplantation of SOD1 KO MuSCs into *mdx* (n=3). (I) *Pgc-1α* gene expression following transplantation of SOD1 KO MuSCs into *mdx* (n=3). Mean ± SD, \**p*<0.05, \*\**p*<0.01 compared to contralateral control for all figures.

To further examine the donor effects of MuSC mitochondrial transfer, we used MuSCs from homozygous *Sod1* knockout mice which exhibit high oxidative damage and mitochondrial myopathy due to deficiency of this important antioxidant enzyme (Jang et al., 2010). In support of previous findings (Ahn et al., 2019; Deepa et al., 2019; Jang et al., 2010; Masser et al., 2016; Muller et al., 2006), significantly reduced state 2 and state 3 oxygen consumption rates were detected in permeabilized fibers from *Sod1-null* mice compared to age-matched WT controls (Figure 4F). Further supporting the notion that MuSC mitochondrial dysfunction drives the myofiber mitochondrial defects, MuSC-specific *Sod1* knockout muscle exhibits high mitochondrial oxidative damage (Figure S5A) and impaired muscle regeneration. Moreover, transplantation of *Sod1-null* MuSCs into *mdx* recipients (Figure 4G) abolished the accretion of functional mitochondria in the host muscle (Figures 4H, 4I), indicating that the bioenergetically-competent donor MuSC mitochondria are vital for the enrichment of oxidative metabolism of the host myofiber. Finally, to demonstrate that the therapeutic potential for MuSC mitochondrial transplantation is not restricted to DMD or dystrophic muscle, we transplanted healthy MuSC mitochondria donors to *Sod1* knockout recipient muscle and found similar boosts in mitochondrial bioenergetic function (Figures S5B, S5C), illustrating the therapeutic potential of MuSC transplantation for treating numerous mitochondrial myopathies.

## Discussion

Here, we demonstrate the consequential role of MuSCs in maintaining myofiber mitochondria homeostasis of DMD skeletal muscle. When MuSC mitochondria are impaired, such as in dystrophic MuSCs, the dysfunctional mitochondria are transmitted to the myofiber during muscle repair. However, the bioenergetic function of dystrophin-deficient muscles can be rejuvenated via transplantation of healthy mitochondria by utilizing donor MuSCs as “delivery vehicles” to ectopically transfer new MuSC-derived mitochondria into host dystrophic myofibers. Despite numerous studies reporting mitochondrial dysfunction in DMD patients and in pre-clinical models, its role in the pathogenesis of DMD has been overlooked and considered a secondary phenomenon to dystrophinopathy (Tabebordbar et al., 2013). Intriguingly, prior to the discovery of dystrophin, DMD was in fact considered a metabolic disorder (Chi et al., 1987; Dreyfus et al., 1954; Timpani et al., 2015a). Nonetheless, a considerable amount of research has focused on restoring dystrophin via gene-or cell-based therapies (Konieczny et al., 2013; Sun et al., 2020; Timpani et al., 2015a), although restoration of mitochondrial function and the metabolic response to MuSC therapies has been largely ignored. Hence, the findings from this study provide insight into the source of the defective mitochondria in dystrophic muscle and validate the ability of MuSCs to revitalize metabolic function of skeletal muscle.

MuSC transplantation will not only enhance bioenergetic function in DMD, but can also restore dystrophin and replenish quiescent MuSC pool in the recipients (Cerletti et al., 2008; Gussoni et al., 1999; Konieczny et al., 2013). Importantly, this approach can be applied in conjunction with recently developed dystrophin-targeting gene therapies (i.e., mini dystrophin, exon skipping using CRISPR/cas) to simultaneously target the genetic etiology of DMD and slow the progression of the disease (Harper et al., 2002; Konieczny et al., 2013; Min et al., 2019; Zhang et al., 2020). However, clinical trials for gene therapy are mostly offered before the onset of severe symptoms and, due to several different mutant variants in the dystrophin gene that confines the pertinence of gene therapy, cell-based therapies may be a better option for many DMD patients (Asher et al., 2020; Tayeb, 2010). Although transplantable MuSC availability may limit the success of translational research, significant breakthroughs in the generation of MuSCs from induced pluripotent stem cells or embryonic stem cells combined with gene correction approaches can improve the clinical outcomes of MuSC-based therapy by amplifying the number of cells transplanted and replenishing the myogenic pool (Chal et al., 2015; Darabi et al., 2012; Min et al., 2019; Mizuno et al., 2010; Pouzet et al., 2001).

Our data demonstrate that the quality of MuSC mitochondria has a profound effect on muscle bioenergetics. Therefore, transplantation of MuSCs to treat mitochondrial myopathy has the potential to be further refined by manipulating donor MuSC mitochondria. For example, endurance-exercising donors enriches the mitochondrial content of MuSCs through PGC-1α-mediated biogenesis (Baar et al., 2002) and primes the cell to readily activate through the mTORC1 pathway (Malam and Cohn, 2014; Rodgers et al., 2014). Alternatively, MuSCs can also be pre-treated or co-delivered with mitochondrial-targeted small molecules or pharmacological agents prior to transplantation to stimulate synthesis of new mitochondria or enhance mitochondrial dynamics in order to optimize functional mitochondrial transfer into the myofiber (Chang et al., 2017; Pauly et al., 2012; Xu et al., 2013). In this context, other mammalian cells such as astrocytes, epithelial cells, and mesenchymal stem cells have been shown to exhibit the ability to transfer mitochondria intercellularly to boost the metabolic function of bioenergetically impaired tissues (Acquistapace et al., 2011; Wang et al., 2011; Wittig et al., 2012). Yet, MuSC-based mitochondrial transfer is unique in that the MuSC-derived myonuclei that regulate mitochondrial homeostasis and myofiber repair also fuse into the regenerating fiber. Thus, for DMD, not only does MuSC transplantation mitigate mitochondrial dysfunction, but may also repair the downstream effects of dystrophin deficiency including, but not limited to, the unstable sarcolemmal membrane, excessive Ca^2+^ influx, and exercise intolerance (de Lateur and Giaconi, 1979; Franco and Lansman, 1990; Hamer et al., 2002), thereby implicating the comprehensive therapeutic benefits of healthy donor MuSCs for dystrophic muscle.

In summary, we report that the oxidative capacity of dystrophic muscle can be influenced by MuSC mitochondria via myogenic fusion. Interestingly, these findings provide a greater understanding of DMD pathophysiology and suggest that mitochondrial dysfunction is an intrinsic feature of the disease, possibly due to accumulation of mtDNA mutations (Wong et al., 2004), that originates in the MuSC and propagates to the myofiber rather than an indirect Ca^2+^-induced damage to myofiber mitochondria (Timpani et al., 2015a). Although this mitochondrial dysfunction debilitates the myogenic program and cell fate mechanisms of dystrophic MuSCs (Chang et al., 2016; Khacho et al., 2016), transplantation of healthy MuSCs resolves the metabolic function of DMD muscle by supplementing the engrafted myofiber with viable myonuclei and mitochondria that reestablish the crosstalk between the mitochondrial- and nuclear-genome. Overall, we provide evidence that mitochondrial transplantation through MuSC fusion revitalizes the mitochondrial function of dystrophic skeletal muscle, unveiling the unexplored potential of an approach that can be used in conjunction with other modes of therapy. This study also illuminates the novel capabilities of the MuSC in muscle metabolism and its therapeutic potential for various other mitochondrial myopathies.

### Limitations of Study

While these findings inform the proof-of-concept effects of MuSC transplantation on mitochondrial function of the myofiber, scaling limitations of donor MuSCs may hinder the therapeutic benefits of this approach for translational research. In addition, we have not tested long-term effects or consequence of multiple injections of donor MuSCs in this study. Furthermore, since every muscle in the body is inflicted in muscular dystrophy, systemic delivery using biomimetic materials will be more appropriate strategy. Nevertheless, despite these limitations, MuSC mitochondria transplantation approach can be still feasible for local mitochondrial myopathy such as hindlimb ischemia, spinal cord injury, volumetric muscle loss, and disuse atrophy and warrants further investigations.

## Methods

### Animals

Mice aged between 2 to 4 months were considered young and mice over 20 months were considered aged. Young and aged *C57Bl/6J* mice used to obtain MuSCs in this study were initially purchased from Jackson Laboratory (Bar Harbor, ME) and maintained in Jang lab mouse colony. *mitoDendra2, H2B-EGFP, Rosa26-tdTomato*, and *DMD-mdx4CV* mice, all with *C57Bl/6J* genetic background, were purchased Jackson Laboratory and bred in our mouse colony. *Sod1 null* mice were initially provided by Dr. Holly Van Remmen (Oklahoma Medical Research Foundation). Female mice were used in this study. All mice were housed, aged, and/or bred in the specific pathogen free condition in Physiological Research Laboratory (PRL) at Georgia Institute of Technology. All procedures were performed in accordance with the approval of the Institutional Animal Care and Use Committee (IACUC) at Georgia Institute of Technology.

### Muscle stem cell (MuSC) isolation and transplantation

MuSCs were isolated through a cell sorting procedure as previously performed.(Cerletti et al., 2008; Han et al., 2018) Briefly, hindlimb muscle tissues were harvested from young (2-4 months of age) and aged (20-24 months of age) mice, and then they were incubated in 20 mL of DMEM media containing 0.2 % collagenase type II (Worthington Biochemical Corporation) and 2.5 U/mL dispase (Gibco) for 90 min. at 37 °C. After tissue digestion, the resulting media were mixed with same volume of stop media (20% FBS in F10), filtered using a cell strainer with a pore size of 70 *μ*m, and then centrifuged (300 *g* for 5 min. at 4 °C) (Allegra X-30R Centrifuge, Beckman Coulter, USA) to obtain the myofiber-associated cell pellet. The cell pellets were washed with Hank’s balanced salt solution (HBSS) containing 2% donor bovine serum (DBS), and the cells were incubated with primary antibodies. For MuSC sorting, a cocktail mixture containing the following antibodies was used: (1) APC conjugated anti-mouse CD11b (1:200; BioLegend), CD31 (1:200; BioLegend), CD45 (1:200; BioLegend), Sca-1 (1:200; BioLegend), and Ter119 (1:200; BioLegend), (2) PE conjugated anti-mouse CD29 (1:100; BioLegend), and (3) biotinylated anti-mouse CD184 (1:100; BD Biosciences). After incubation for 30 min. at 4 °C, the primary antibodies-treated cells were washed, centrifuged (300 *g* for 5 min. at 4 °C), and then treated with a secondary antibody (Streptavidin PE-Cy7) (1:50; Invitrogen) for 20 min. at 4 °C. Following propidium iodine (PI) treatment and strainer filtration (70 *μ*m), the MuSCs (PI, CD11b, CD45, Sca-1, and Ter119; negative selection, CD29 and CD184; positive selection) were subsequently isolated by a Fluorescence activated cell sorting (FACS) (BD FACS Aria III, BD Biosciences, USA). 24 hours after BaCl_2_ (1.2% w/v) injury to the tibialis anterior muscle (TA) of the recipient mouse to induce regeneration, 200,000 freshly sorted MuSCs were injected into the TA intramuscularly. 4 weeks following MuSC transplantation, mice were euthanized and the TAs were dissected for tissue analysis. Sham injury with BaCl_2_ injection but no MuSC transplantation was used as the contralateral control. No differences in mitochondrial content or myofiber respiration were observed between naive *mdx* muscle and BaCl_2_-injured muscle 28 days after injection.

### β1-integrin and CXCR4 cluster analysis

Using β1-integrin (CD29) and CXCR4 (CD184) as stemness markers of MuSCs from FACS sorting data, we clustered purified MuSCs from wildtype and *mdx* mice based on CD29 and CD184 expression using SPADE tree analysis to delineate the heterogeneity of the MuSC population. Each node represents a set of MuSCs clustered together based on expression of all surface markers (CD11b, CD45, Sca-1, Ter119, CD29, and CD184). Within the SPADE tree, nodes were then manually grouped to congregate nodes with high CD29/high CD184 expression, high CD29/low CD184 expression, low CD29/high CD184 expression, and low CD29/low CD184 expression. The total number of cells within each of these groups were then counted and compared between wildtype and *mdx* MuSC populations.

### Muscle stem cell culture and staining

Following MuSC isolation, 8,000 purified cells were seeded into an ibidi 96 well glass bottom plate in F10 media supplemented with 20% horse serum, 1% glutaMAX, 1% penicillin/streptomycin, and 25 ng/mL basic fibroblast growth factor (bFGF). For quiescent MuSCs, cells were fixed with 4% paraformaldehyde after 12 hours to allow cells to adhere to bottom of the plate. To proliferate MuSCs while inhibiting differentiation, 25 ng/mL bFGF was supplemented daily. To differentiate MuSCs and facilitate fusion into myotubes, MuSCs were first treated with 25 ng/mL bFGF for 4 days and then allowed to differentiate and fuse for another 6 days before fixation. Fixed cells were then stained by treating cells with Hoechst 33342 (1:1000 dilution) for DAPI and phalloidin-iFluor 555 (1:500 dilution) for F-actin, diluted in blocking buffer (2% BSA, 0.5% goat serum, 0.5% Triton X-100 in PBS). After staining, z-stack images of cells were taken on a Zeiss 700 Laser Scanning Confocal microscope. Mitochondrial content was quantified by measuring relative fluorescent intensity of *Dendra2* signal in maximum intensity projections.

### Mitochondrial respiration of muscle stem cells

Using the Lucid Scientific RESIPHER device on a 96-well plate, oxygen flux of the cell monolayer can be quantified real-time throughout MuSC culture and myogenesis. Oxygen flux was quantified in wells with 8,000 seeded MuSCs during the first 4 days with 25 ng/mL bFGF supplementation to proliferate MuSCs and during the next 6 days without bFGF to allow MuSC differentiation and fusion. For comprehensive analysis of mitochondrial respiration in MuSCs, 8,000 cells per well in a Seahorse XFp miniplate were supplemented with 25 ng/mL for 8 days until confluent. Prior to running the Seahorse assay, media was changed to XF Base DMEM with 1 mM pyruvate, 2 mM glutamine, and 10 mM glucose. Sensor cartridges were loaded for sequential injections to result in final concentrations of 1 µM oligomycin, 1 µM carbonyl cyanide 3-chlorophenylhydrazone, and 0.5 µM rotenone. MuSCs and the cartridge were placed in a Seahorse XFp Analyzer to test mitochondrial function at various stress conditions.

### Cross-sectional and single fiber imaging

Following euthanasia, mouse hindlimbs were fixed in 4% PFA and the tibialis anterior muscles were dissected. For cross-sectional images, muscles were prepared as previously described (Anderson et al., 2019) where muscles were cryopreserved with 20% sucrose overnight then frozen in liquid nitrogen-cooled 2-methylbutane. Muscles were sectioned at 10 µm in a cryostat and stained with phalloidin (for F-actin) (Thermo A22287) and DAPI (Vector Labs H-1200) for fluorescent imaging on a Zeiss 700 Laser Scanning Confocal microscope. For single fiber images, fixed muscles were first stained with phalloidin and DAPI, then mechanically separated into single muscle fibers as previously described (Mohiuddin et al., 2019). Isolated muscle fibers were mounted on slides for imaging on a confocal microscope.

### Western blot analysis

Tibialis anterior muscles were homogenized in BioMasher homogenization tubes (VWR KT749625-0030) in RIPA lysis buffer (VWR 97063-270) supplemented with Roche cOmplete Mini Protease Inhibitors (Roche 04693124001) and PhosSTOP Phosphatase Inhibitors (Roche 04906837001) at 1 mL lysis buffer per 100 mg muscle tissue. After 3 freeze-thaw cycles, muscle homogenates were centrifuged at 18,400 *g* for 10 minutes and the supernatants were collected. The supernatants were normalized for protein concentration using a BCA assay kit (Thermo 23225) and mixed with Laemmli buffer (Bio-Rad 161-0737). 50 µg of the protein mix were run through a 4-20% Criterion TGX gel (Bio-Rad 5671093) at 150 V for 85 minutes and transferred to a PVDF membrane using the Trans-Blot Turbo System at 2.5 A for 7 minutes. Following antibody staining, membranes were imaged on Li-Cor Odyssey CLx-1050 Infrared Imaging System and bands were quantified on Li-Cor Image Studio V5.2. Membranes were subsequently stained with Ponceau solution (Sigma P7170) as a loading control and imaged on Amersham.

### RNA isolation and quantitative polymerase chain reaction

Muscles were homogenized in a similar method to Western blot preparation with the exception of RLT buffer supplied in the RNeasy® Mini Kit (Qiagen 74104) supplemented with 1% β-mercaptoethanol (Sigma M6250) instead of RIPA lysis buffer. Following collection of the supernatant, RNA was isolated according to the protocol provided by Qiagen RNeasy® kit. RNA concentration was measured in a NanoDrop One and samples were normalized according to RNA content. The Applied Biosystems High Capacity cDNA Reverse Transcription Kit (4368814) was used to reverse transcribe the isolated RNA into cDNA. The cDNA was combined with Applied Biosystems PowerUp SYBR Green Master Mix (A25742) and corresponding primers in 96-well plates and run in an Applied Biosystems StepOnePlus Real-Time PCR system to perform the qPCR reactions. β-actin and B2m, which we found to be stably expressed in our muscle samples, were used as housekeeping genes to quantify relative fold induction. For the PCR array data, cDNA was mixed with PowerUp SYBR Green Master Mix, loaded into the Qiagen Mouse Mitochondria RT^2^ Profiler PCR Array (330231), and run in an Applied Biosystems StepOnePlus to quantify the data.

### mRNA sequencing library preparation

mRNA isolation and preparation were conducted as previously described (Aguilar et al., 2016). Briefly, sorted MuSCs were collected in Trizol (Thermo) and total RNA was purified with miRNeasy Micro kit (Qiagen) following the manufacturer’s instruction. The quality of RNA and its concentration were measured by a spectrophotometer (Nanodrop 2000c) and Bioanalyzer (Agilent 2100). cDNA libraries were produced using SmartSeq4 protocol (Clontech) using 10 ng of total RNA following the manufacturer’s instructions. We sequenced individual library using 76 paired-end reads with an average depth of 50M reads per sample.

### Analysis of differentially expressed gene

RNA sequencing reads were first processed with Trimmomatics (ver. 0.38) (Bolger et al., 2014) to remove potential adapter sequences and to trim low quality sequences. Trimmed RNA-Seq reads were then mapped to Mus musculus genome (GRCm38) and gene expression is quantified using RSEM (ver. 1.2.28) (Li and Dewey, 2011). We used tximport (ver. 1.8) to import expected read counts into DESeq2 (ver. 1.2) Bioconductor package (Love et al., 2014). We removed genes with FPKM < 1 across all samples to exclude low expressed genes. Differential gene expression analysis was performed using DESeq2. Genes were determined to be significantly differentially expressed if an absolute average expression fold change between WT and mdx groups is greater than 2 with P-value below 0.01 after multiple correction. Gene Set Enrichment Analysis (GSEA) was performed to determine significantly enriched GO terms and pathways (Subramanian et al., 2005).

### Permeabilization of muscle fibers

Immediately following euthanasia and dissection of the tibialis anterior, fresh muscles were places in cold buffer X (7.23 mM K_2_EGTA, 2.77 Ca K_2_EGTA, 20 mM imidazole, 20 mM taurine, 5.7 mM ATP, 14.3 mM phosphocreatine, 6.56 mM MgCl_2_-6H_2_O, 50 mM K-MES) as previously described (Berru et al., 2019). Fiber bundles with approximate wet weights of 4 mg were mechanically separated under a dissecting scope with fine forceps and permeabilized with 30 µg/ml saponin for 40 minutes. For transplanted muscles, a NIGHTSEA Stereo Microscope Fluorescence Adapter (Electron Microscopy Sciences SFA-LFS-RB, SFA-LFS-GR) was used to locate and isolate engrafted fibers. Following permeabilization, fiber bundles were washed in cold buffer Z (105 mM K-MES, 30 mM KCl, 10 mM KH_2_PO_4_, 5 mM MgCl_2_-6H_2_O, 0.5 mg/ml BSA, 1 mM EGTA) for 15 minutes to remove any residual saponin.

### Mitochondrial respiration of myofibers

Following permeabilization of fiber bundles, samples were loaded into an Oroboros O2k oxygraph with 2 mL buffer Z without MgCl_2_ and supplemented with 2.5 mM Magnesium Green (Thermo M3733) in order to simultaneously measure oxygen consumption and relative ATP production rates. Prior to loading samples in the chamber, the system was calibrated using various titrations of ATP to ADP ratios to determine relative ATP production due to the higher binding affinity of Magnesium Green to ATP compared to ADP. After loading samples, the chamber was equilibrated for 15 minutes to reach basal (State 1) respiration rates. Glutamate (10 mM) and malate (2 mM) were then added to the chambers for complex I-linked State 2 respiration rates, followed by the addition of ADP (5 mM) to initiate State 3 respiration. Oxygen consumption rates were expressed as pmol/sec/mg fiber wet weight while relative ATP production rate was measured as changes in the ATP to ADP ratio. All measurements were conducted at 37 °C and a working [O_2_] range ∼250 to 175 µM.

### Endurance exercise

Endurance-exercised donor mice were run on a 4-lane rodent motor-driven treadmill (Omnitech, USA). For proper adaptation to the treadmill, mice were pre-trained for one week at 5-10 meters/minute for 10-20 minutes each day. Following pre-training, each training session for the main exercise was progressively increased from 5-10 m/min to 15-20 m/min over 50-60 minutes each day. The mice underwent a warm-up session (2-3 m/min) and a cool-down session (3-5 m/min) before and after the main exercise, respectively. These exercises continued for 8 weeks at 5 sessions each week. After completion of the exercise protocol, MuSCs were purified from exercised donor mice one week after the last session to allow metabolic changes of the MuSC and skeletal muscle to occur.

### Statistical Analyses

All statistical analyses in this study were performed on GraphPad Prism 7 and data is presented as mean ± standard deviation (SD). Samples sizes were chosen using G*Power based on preliminary experiments to ensure adequate statistical power. Normality of data was tested with the Shapiro-Wilk test. For all experiments comparing wildtype data to *mdx* data and *mdx* data to transplant data, unpaired two-tailed t-tests were used to determine statistical difference. A *p*-value of less than 0.05 was considered statistically significant.

### Accession codes

RNA-Seq data from this study has been archived at the NCBI Sequence Read Archive under BioProject [PRJNA:625722]

## Acknowledgements

Research reported in this publication was supported by the National Institute of Arthritis and Musculoskeletal and Skin Diseases of the National Institutes of Health under Award Number R21AR072287 (YCJ), S&R Foundation, and grants from the Parker H. Petit Institute of Bioengineering and Bioscience, and the Center for Regenerative Engineering and Medicine. The content is solely the responsibility of the authors and does not necessarily represent the official views of the National Institutes of Health. We acknowledge the staffs at the Children’s Healthcare of Atlanta for valuable discussions and insight. We also thank the Physiological Research Laboratory and core facilities at the Parker H. Petit Institute of Bioengineering and Bioscience at the Georgia Institute of Technology for the use of shared equipment, services, and expertise.

## Author Contributions Statement

M.M., J.J.C., and Y.C.J. designed the experiments, analyzed the data, and wrote the manuscript. M.M., J.J.C., N.H.L., H.J., S.E.A., W.M.H., T.Z.P., and B.A. conducted the experiments, contributed to methodology, and analyzed the data. A.J.G. and C.A.A. provided significant contributions to methodology and conceptual justification. N.H.L. and Y.C.J. generated and maintained animals used in this study. All authors reviewed the manuscript.

## Supplemental Information

**Supplemental Figure S1.**
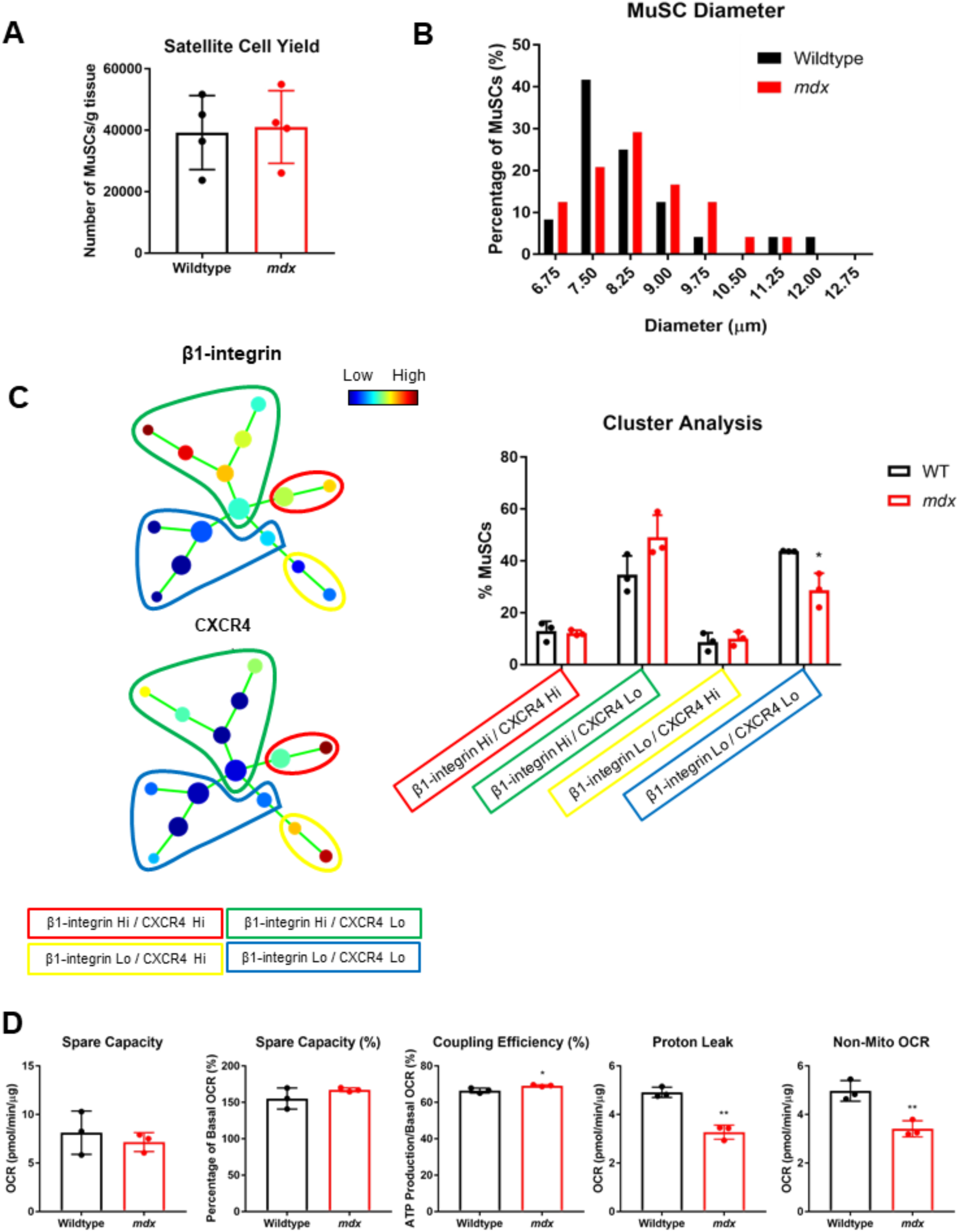
Dystrophic muscle stem cells have impaired mitochondrial function and myogenic potential. (A) MuSC yield per gram of tissue from wildtype and *mdx* skeletal muscle (n=4). (B) Histogram of diameter of 25 MuSCs from mitoDendra2 and *mdx*/mitoDendra2 muscle. (C) Cluster analysis of wildtype and *mdx* MuSCs using SPADE tree analysis (n=3). (D) Quantifications of oxygen consumption rate using Seahorse XF flux analyzer (n=3).

**Supplsemental Figure S2.**
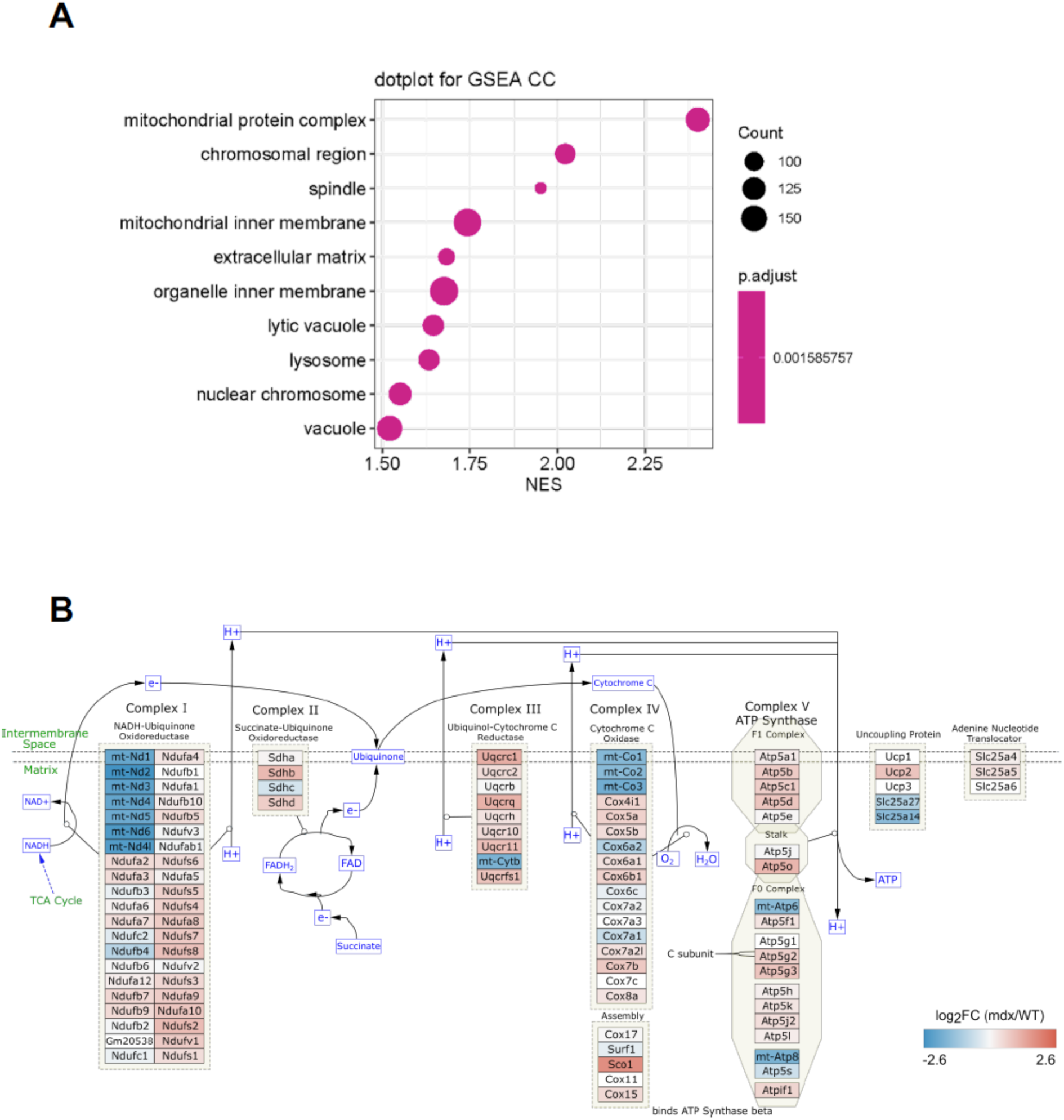
Dystrophic muscle stem cells upregulate mitochondrial gene expression. (A) Gene set enrichment pathway analysis of *mdx* MuSCs compared to wildtype MuSCs. (B) Diagram and changes in gene expression of mitochondrial electron transport chain and oxidative phosphorylation using RNA seq. data.

**Supplemental Figure S3.**
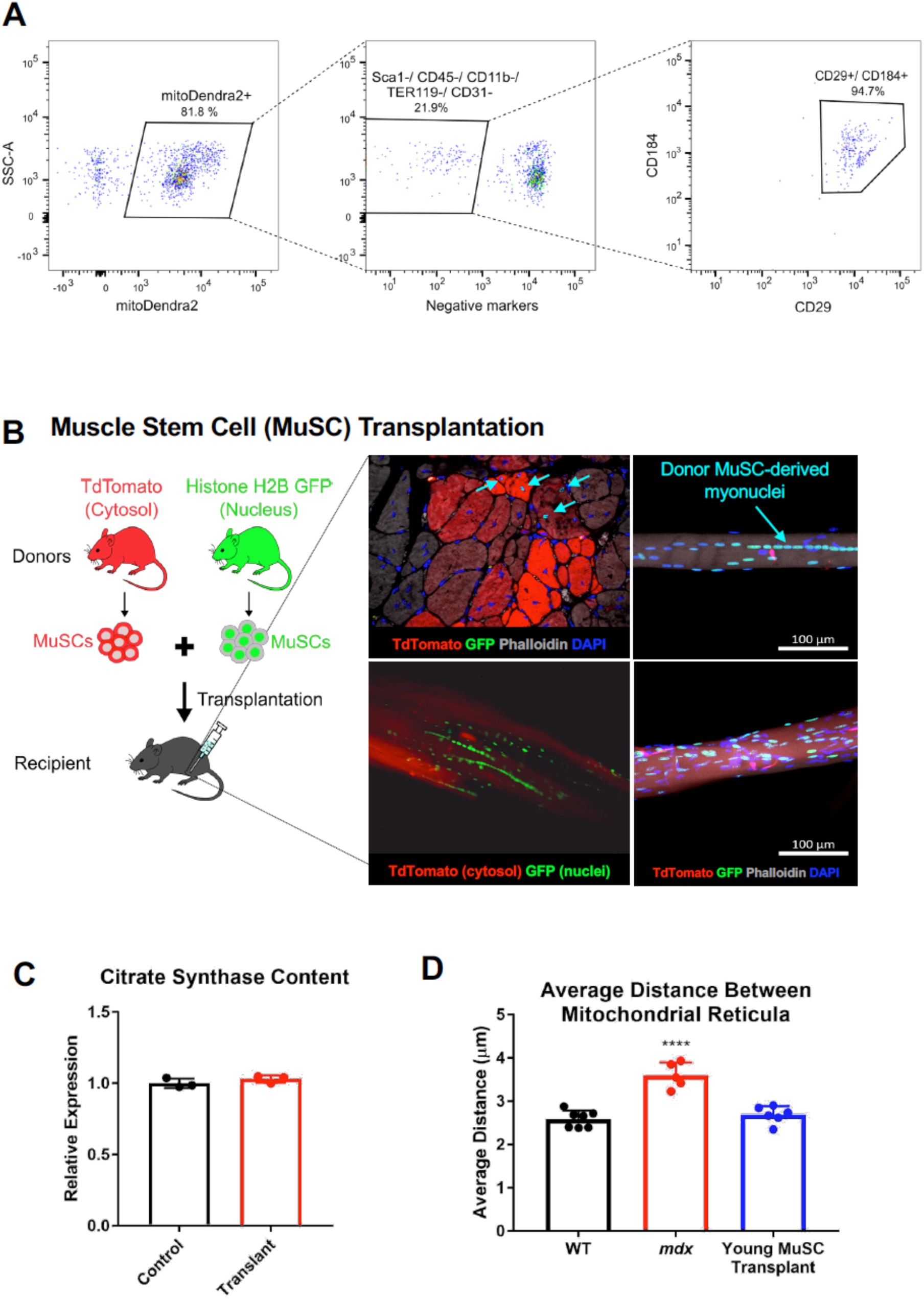
Purification and transplantation of quiescent muscle stem cells. (A) Representative gating of quiescent MuSCs using fluorescence-activated cell sorting. (B) Schematic for transplantation of tdTomato and H2B-EGFP MuSCs into *mdx* TA. Cross-sectional and single fiber images for transplanted muscle. tdTomato in red, H2B in green, actin in gray, myonuclei in blue. (C) Citrate synthase Western blot following transplantation of young MuSCs into *mdx* (n=3). (D) Mitochondrial content (relative fluorescent units) along 10 µm of fiber length. \**p*<0.05, \*\**p*<0.01, \*\*\*\**p*<0.0001 compared to wildtype for all figures.

**Supplemental Figure S4.**
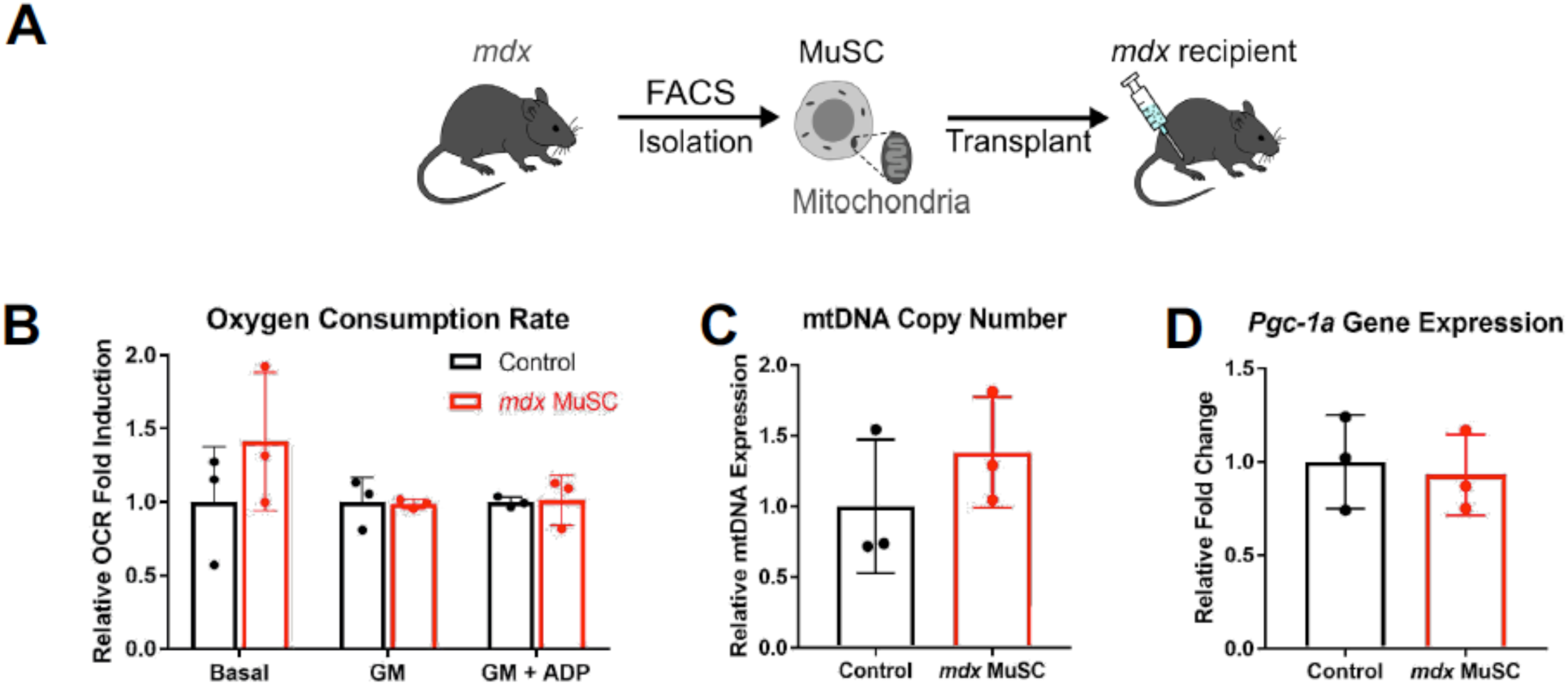
Transplantation of dystrophic muscle stem cells into dystrophic recipient. (A) Schematic diagram of transplantation of *mdx* MuSC donor into *mdx* host. (C) OCR of permeabilized fibers following transplantation of *mdx* MuSCs into *mdx* host (n=3). (D) mtDNA copy number following transplantation of *mdx* MuSCs into *mdx* host (n=3). (E) *Pgc-1α* gene expression following transplantation of *mdx* MuSCs into *mdx* recipients (n=3).

**Supplemental Figure S5.**
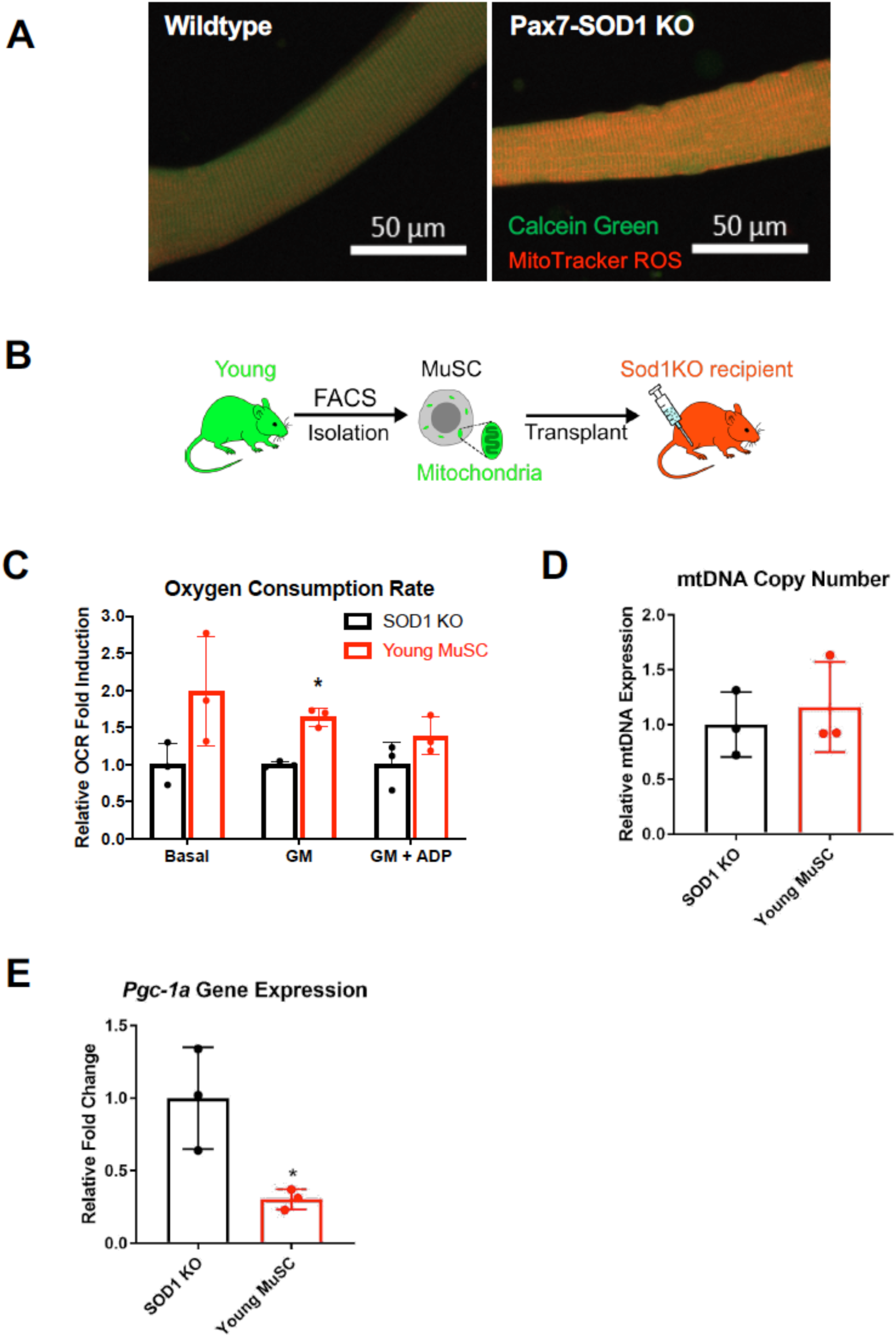
Transplantation using an SOD1 knockout model. (A) Confocal images of single fibers from FDB of wildtype and Pax7-SOD1 knockout mice stained with Calcein green and Mitotracker CM-H_2_XROS. (B) Schematic for transplantation of young MuSCs into SOD1 KO TA. (C) OCR of permeabilized fibers following transplantation of young MuSCs into SOD1 KO (n=3). (D) mtDNA copy number following transplantation of young MuSCs into SOD1 KO (n=3). (E) *Pgc-1α* gene expression following transplantation of young MuSCs into SOD1 KO (n=3). \**p*<0.05 compared to contralateral control for all figures.

